# A multi-level society comprised of one-male and multi-male core units in an African colobine (*Colobus angolensis ruwenzorii*)

**DOI:** 10.1101/641746

**Authors:** Samantha M. Stead, Julie A. Teichroeb

## Abstract

A few mammalian species exhibit complex, nested social organizations, termed multi-level societies. Among nonhuman primates, multi-level societies have been confirmed in several African papionin and Asian colobine species. Using data on individually-recognized Rwenzori Angolan colobus at Lake Nabugabo, Uganda, we document the first multi-level society in an African colobine. The study band comprised up to 135 individuals living in 12 socially and spatially distinct core units that ranged in size from 4 to 23 individuals. These core units shared a home range, and fissioned and fused throughout the day. Using the association indices between core units, we employed hierarchical cluster analyses and permutation tests to show that some core units clustered into clans. Thus, we confirm three tiers of social organization for Rwenzori Angolan colobus: core unit, clan, and band. The social organization of this subspecies is unlike any reported previously in a nonhuman primate, with about half the core units containing a single adult male and the others containing multiple reproductive adult males. Preliminary data show males to transfer within the band and female to transfer outside of the band, which suggests that, like Hamadryas baboons, this subspecies could provide insight into the selective pressures underlying hominin social organization.

## Introduction

### Description of multi-level societies

Decades of research have been devoted to understanding the underlying components of animal social systems (mating systems, social organization, and social structure)^1–2^. Social organization, a description of group size, composition and cohesion, is known to vary both across and within species, reflecting local ecological and social conditions^3–7^. Social systems that display short-term changes in one or more of these three parameters (size, composition, and cohesion) are said to exhibit fission-fusion dynamics^8^. In some species, fission and fusion occurs predictably along certain social boundaries. These species are said to live in multi-level (or modular) societies, which are complex social systems made up of basic social units (hereafter referred to as core units) that fission and fuse with one another in a hierarchical manner^9–10^. Core units are socially and often spatially distinct, with direct social interactions occurring most frequently between individuals in the same core unit^9^. This hierarchy of social and spatial association can extend to multiple tiers, with more complex systems showing up to four tiers of non-random association (e.g., geladas^11^, hamadryas baboons^12^, elephants^13^).

Multi-level societies have been described in several mammalian species, including African elephants (*Loxodonta africana*)^13–14^, giraffes (*Giraffa camelopardalis*)^15^, plains zebras (*Equus burchelli*)^16^, prairie dogs (*Cynomys ludovicianus*)^17^, khulans (*Equus hemionus*)^18^, sperm whales (*Physeter microcephalus*)^19^, and bottlenose dolphins (*Tursiops* sp.)^20^. Within the primate order, multi-level societies have been confirmed in Asian colobines (*Rhinopithecus* spp.^21–22^, *Nasalis larvatus* ^23^), African papionins (*Theropithecus gelada*^11^, *Papio hamadryas*^24^, *Papio papio*^25–27^), and humans^28–31^. Additionally, it has been suggested that uakaris (*Cacajao calvus ucayalii*)^32^ and drills (*Mandrillus leucophaeus*)^33^ live in multi-level societies; however, more conclusive data are needed.

Multi-level societies of non-human primates show exceptional group sizes, with aggregations of several hundred individuals observed at times (reviewed by Grueter and colleagues^34^). Core units are known to be one-male/multi-female (OMU) and all-male (AMU), with no stable multi-male/multi-female core units (MMU) documented thus far^11, 21, 23^ ^24^, although see Bowler and colleagues^32^ and Cui and colleagues^35^ for some weak evidence in uakaris and an Asian colobine (*Rhinopithecus bieti*), respectively. In African papionins, ‘follower’ or ‘secondary’ males can associate regularly with one or more OMU^25,36–37^; however, these males rarely have sexual access to females. Up to four tiers of non-random association have been documented in African papionins (*Theropithecus gelada*^11^; *Papio hamadryas*^24^, *Papio papio*^25^) and up to three tiers in Asian colobines (*Nasalis larvatus*^23^, *Rhinopithecus* ssp.^21,38^).

### Evolution of multi-level societies

There are many costs and benefits of group living, mostly related to food availability^3–6^, predator threat^39–40^ and conspecific threat^7,41^. These factors can vary as groups move through time and space, collectively influencing optimal group size. As such, the ability to modify group size in response to fluctuating ecological and social conditions has obvious fitness benefits, which are thought to have driven the evolution of multi-level societies. The bachelor threat hypothesis proposes that core units in multi-level societies associate so that resident males can collectively deter bachelor male(s)^16,38,42–44^. For example, gelada males from different OMUs associate more closely in the presence of bachelor males, which, in geladas, form AMUs that far outnumber the resident male of a unit^45^. Predation threat is another selective pressure that is thought to have driven the evolution of some tiers in multi-level societies. For instance, in hamadryas baboons, on mornings after predator calls were heard, bands remained more cohesive while foraging than they typically do^46^. This grouping behaviour is thought to counter predation threat through risk dilution and increased detection rates^39,40,47^. In the absence of predator calls, hamadryas bands tend to break up into sub-units to forage, as this reduces the costs of scramble competition for food^46^. In fact, in several primate multi-level societies, fissioning occurs in response to low food availability as a way to reduce competition. For instance, during periods of low food availability, hamadryas baboon bands fission more frequently into OMUs^24,46^ and golden snub-nosed monkey troops fission into herds (composed of OMUs and AMUs)^38^. Thus, there is evidence from several species that flexible grouping in multi-level societies evolved to allows animals to optimize their group size in response to fluctuating conditions (i.e., bachelor threat, predator threat, food availability).

### Rwenzori Angolan colobus

To better understand the evolution of multi-level societies, it is important to first document the presence of these social systems across the animal kingdom. For decades, Rwenzori Angolan colobus (also referred to as Adolf Friedrich’s Angolan colobus) have been known to differ from their congeners. Early research on a population in the Nyungwe Forest in Rwanda found exceptionally large groups, with more recent data showing over 500 individuals travelling together^48–50^. These group sizes are extreme outliers among black-and-white colobus (*Colobus* spp.) where groups typically average less than 20 individuals (reviewed by Fashing^51^). Rwenzori Angolan colobus at our study site at Lake Nabugabo, Uganda form large groups of about 200 individuals and exhibit a greater degree of fission-fusion dynamics than reported at Nyungwe^52^. The aims of this study were 1) to determine if Rwenzori Angolan colobus at Nabugabo are forming a multi-level society and if so, 2) to determine the core unit composition and tier number.

## Materials and methods

### Study site

This study was conducted at Lake Nabugabo, Masaka District, central Uganda (0°22’-12°S and 31°54’E). Nabugabo is a small lake (8.2 × 5 km) that lies just to the west of Lake Victoria at an elevation of 1,136 m. Rwenzori Angolan colobus (*Colobus angolensis ruwenzorii*) occupy patches of lowland moist evergreen forest around Lake Nabugabo. We studied this subspecies in and around the Manwa Forest Reserve (~280 ha), located near the trading centers of Bukumbula and Bbaale^53^.

### Study population and data collection

JAT began habituating a large band (TR band) of Rwenzori Angolan colobus in September 2013 and follows have been conducted several days per month in most months since that time. Here, we used data collected from August 2017 to August 2018 by SMS and two field assistants where TR band was followed a mean of 11.23 days per month (range: 4-25) from approximately 7:30 am to 4:30 pm (*N* = 150 days). TR band consisted of 119-135 colobus during this period, organized into 12 core units that range in size from 4 to 23 individuals.

Throughout the study period, one core unit was followed each observation day, with an effort made to rotate between each of the 12 core units throughout the month. During contact time with the colobus, we conducted focal-time follows, where one individual was followed for two hours and a scan was taken on them every 15-minutes, which included a GPS point. With this data collection regime, we were able to recognize all individuals in TR band by January 2018 based on various attributes, including tail shape, broken fingers, nipple colours, and eyebrow curvature. Of the GPS points recorded during focal-time follows, here we report spatial data using two points per day to ensure independence – the point recorded in the morning upon contact with the core unit and the one recorded at the end of the observation day.

Scan sampling was used to record association patterns between core units^54^. Every two hours we recorded the identity of all core units that were within a 50-meter radius of the focal core unit. This two-hour time period was chosen to capture the frequent fissioning and fusing of core units while also allowing enough time for associations among core units to change. A distance of 50 metres was selected as an appropriate cut-off to record two or more core units as being ‘in association’ due to three main factors: 1) Since core units were, at times, observed to be over 500 metres apart, 50 metres seemed to be a reasonable distance at which we could be sure that core units were intentionally moving into or staying in proximity of one another. 2) This distance is commonly used to define the occurrence of intergroup encounters in primates, which involve two groups coming close enough to see one another and interact^55–57^ 3) In the dense forest that our study species occupies, it was difficult to positively identify core units that were clustered beyond the units in direct association of the focal unit, without leaving the focal unit.

### Data analyses

We describe the composition of core units and the association patterns of *C. a. ruwenzorii*. Two tiers of social organization (the core unit and band) were detected by observing individual interactions, analyzing associations between core units and plotting spatial data. Following other researchers that have assessed the number of tiers present in multi-level societies^11,13^, we used a hierarchical cluster analysis to determine whether a third tier of organization (the clan) was present for Rwenzori Angolan colobus by examining whether some core units associated preferentially with one another. We used the scans of core unit association (*N* = 544) to calculate association indices between units. The association index that we chose was the simple ratio association index (AI) because we had good visibility during scans, and we could positively identify all core units in association with the focal unit^58^. This ratio was calculated as AI = *N*_AB_/(*N*_A_ + *N*_B_) or the number of times that two units were in association divided by the total number of scans in which either core unit was present. We assessed four different clustering methods to determine which had the best fit with the data (average linkage, Ward’s weighted, complete linkage, and single linkage). The average linkage method led to the highest cophenetic correlation coefficient (CCC = 0.812), indicating that the dendrogram it produced had the best fit with the data^59^, so we used this dendrogram for the rest of the analyses. To determine if there was preferential association among some core units, we counted the number of bifurcations present in the average linkage dendrogram and graphed the cumulative bifurcations to identify “knots” where the slopes below and above knots differed^60^. We assessed the significance of the slope change at knots using the Wilcoxon two-sample Z test^13^.

Further, we used permutation tests for preferred/avoided companionships^61^ between our core units to test the null hypothesis that core unit association occurred at random. We permuted matrices of association indices and our test statistics were the CV (i.e., coefficient of variations) of the association indices (to indicate preferred associations) and the proportion of non-zero elements (to indicate avoidance between some units)^60^. We permuted matrices 1000, 2000, 5000, and 10,000 times until the p-values stabilized and we present the results from the run of 10,000 random permutations. We performed cluster analysis and permutation tests in SOCPROG v2.8^59^ and alpha values less than 0.05 were deemed significant.

### Ethics approval

This research was done with the permission of the Uganda Wildlife Authority (Permit #: UWA/TDO/33/02) and the Uganda National Council of Science and Technology (Permit #: NS537). All methods were carried out in accordance with guidelines and regulations provided by the University of Toronto Animal Care Committee (Approved Protocol #: 20011416).

## Results

We identified a total of 135 individuals and followed them from August 2017 to August 2018. We found that these individuals did not associate randomly with one another but rather in three hierarchical tiers. The terminology that we use to describe these tiers is consistent with existing papers that have studied multi-level societies in Asian colobine species^38^. Previous research on Asian colobines has not described a social tier comparable to the clan and so we adopted terminology from research on African papionins^24^.

### Core units

The lowest tier of non-random association was identified by observing individual-level proximity and behaviour. Individual were consistently surrounded by the same nearest neighbours, all of which rested and moved together throughout the day. These groups of individuals were referred to as ‘core units’. Individuals within a core unit were not only spatially cohesive, but also socially cohesive; direct interactions (e.g., grooming^52^, copulations) were only ever observed to occur between members of the same core unit. Note that all adult males in multi-male/multi-female core units were seen to mate with females, with the exception of one seemingly very old male in core unit PS, who sometimes trailed his unit.

Based on these criteria, we identified 12 core units, which ranged in size from 4 to 23 individuals. The number of adult males per core unit ranged from 1 to 8 and the number of adult females ranged from 1 to 6. At the end of the study period, the 12 core units had a mean of 11.25 individuals per core unit, 2.67 adult males per core unit and 3.75 adult females per core unit. Four core units (AL, FA, MA, PO) were consistently uni-male/multi-female throughout the study period, five units were consistently multi-male/multi-female (AN, LI, LO, NE, PS), and three units changed from uni-male to multi-male or vice versa (BR, FU, PH) (Table 1).

**Table 1.**
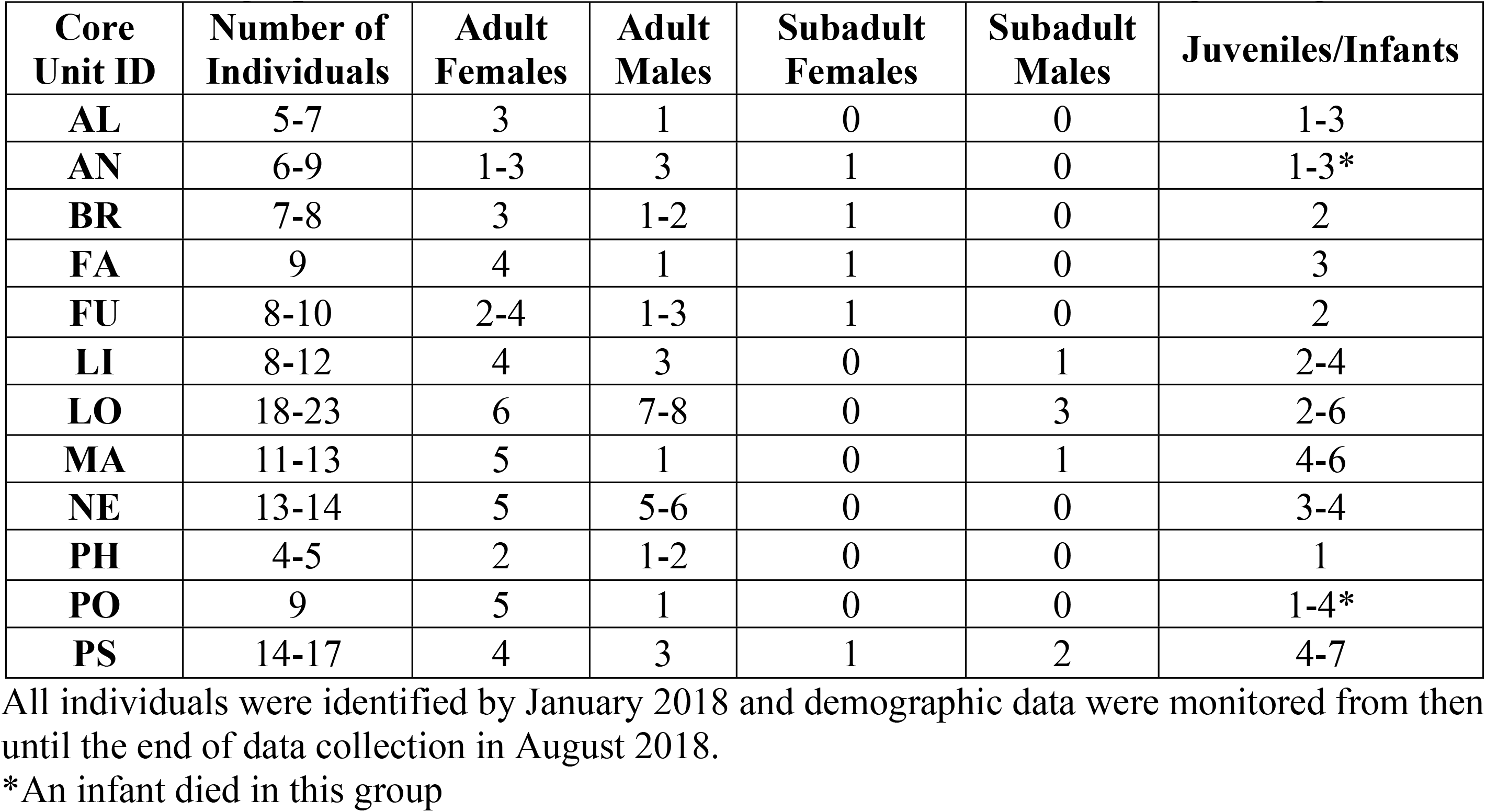
Demographic Data for 12 Core Units of *C. a. ruwenzorii* at Nabugabo, Uganda.

| Core Unit ID | Number of Individuals | Adult Females | Adult Males | Subadult Females | Subadult Males | Juveniles/Infants |
| --- | --- | --- | --- | --- | --- | --- |
| AL | 5-7 | 3 | 1 | 0 | 0 | 1-3 |
| AN | 6-9 | 1-3 | 3 | 1 | 0 | 1-3* |
| BR | 7-8 | 3 | 1-2 | 1 | 0 | 2 |
| FA | 9 | 4 | 1 | 1 | 0 | 3 |
| FU | 8-10 | 2-4 | 1-3 | 1 | 0 | 2 |
| LI | 8-12 | 4 | 3 | 0 | 1 | 2-4 |
| LO | 18-23 | 6 | 7-8 | 0 | 3 | 2-6 |
| MA | 11-13 | 5 | 1 | 0 | 1 | 4-6 |
| NE | 13-14 | 5 | 5-6 | 0 | 0 | 3-4 |
| PH | 4-5 | 2 | 1-2 | 0 | 0 | 1 |
| PO | 9 | 5 | 1 | 0 | 0 | 1-4* |
| PS | 14-17 | 4 | 3 | 1 | 2 | 4-7 |

All individuals were identified by January 2018 and demographic data were monitored from then until the end of data collection in August 2018.
\*An infant died in this group

Individuals from different core units were spatially separate from one another, usually occupying different, but neighboring, tree crowns. At times, multiple core units were observed to occupy the crown of large trees; however, they always remained spatially distinct. Core units were usually tolerant of one another and most social interactions that did occur between units were indirect (i.e., vocalization, displaying) and typically benign, as is shown in other primate multi-level societies^34^. The 12 core units were not territorial with one another and shared a home range of about 1.5 km^2^ throughout the study period (Fig 1). This home range size was estimated using GPS data collected for core units at the beginning and end of each focal follow. If these 12 core units simply share a home range, then we would expect them to encounter one another randomly. However, if Rwenzori Angolan colobus form a multi-level society, and thus coordinate their activity with one another, we expect to see delineated patterns of association, which we show in the next section.

**Fig 1.**
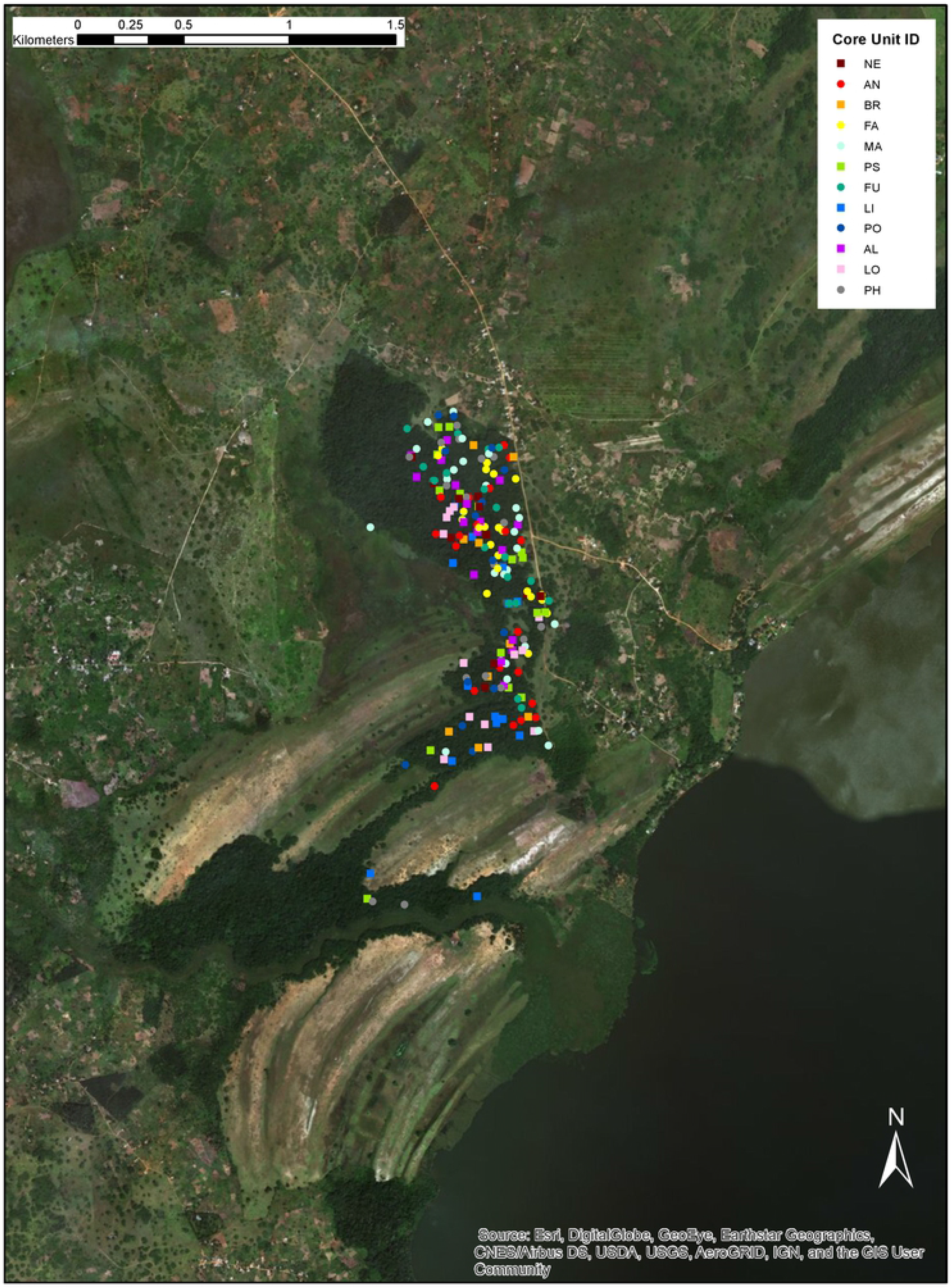
**Spatial Data** GPS points (*N* = 207) collected at the beginning and end of every observation day on a focal unit; the first GPS point of that day marks where the focal core unit was first located and the second marks where the focal core unit ended up. GPS data was collected from July 14^th^, 2017 to May 16^th^, 2018. Each of the core units (*N* = 12) are represented by a different coloured point. Map created using ArcGIS.

### Two clans and one band

We found that the focal core unit was usually in association with (i.e., within 50 m of) at least one of the other 12 identified core unit(s), which we collectively refer to as TR band. A ‘band’ comprises multiple core units that share a home range and maintain or tolerate proximity with one another. Over 150 observation days, there were only 3 days where another core unit(s) from TR band was not within 50 m of the focal core unit during scans. Thus, the focal core units were in association with another unit(s) on at least 98% of observation days. The mean number of core units in association with the focal core unit was 2.75 (±0.51 SD, range: 0-9). The mean number of scans where no other core unit was in association with the focal core unit was 10.9 (±7.4 SD), for an average estimated 22.56% of time spent out of association. In other words, focal core units were within 50 m of one or more core unit(s) from TR band in 77.44% of association scans despite having an overlapping home range of at least 1.5 km^2^ (Fig 1). This suggests that core units from TR band are not randomly encountering one another due to their shared home range but rather, that they are intentionally maintaining proximity with one another. This is further supported by the fact that we regularly observed associations between core units to last over several 2-hour scans, sometimes into the next day, and core units often moved together. Lagged association rates, which were defined as the probability that two core units were associated given their association in the previous scan, plotted against null association rates show that there was a tendency for units to stay together over several 2-hour scans (Fig 2). Since we followed one core unit per day, our data collection regime did not allow us to determine if certain core units stayed together over even longer time periods.

**Fig 2.**
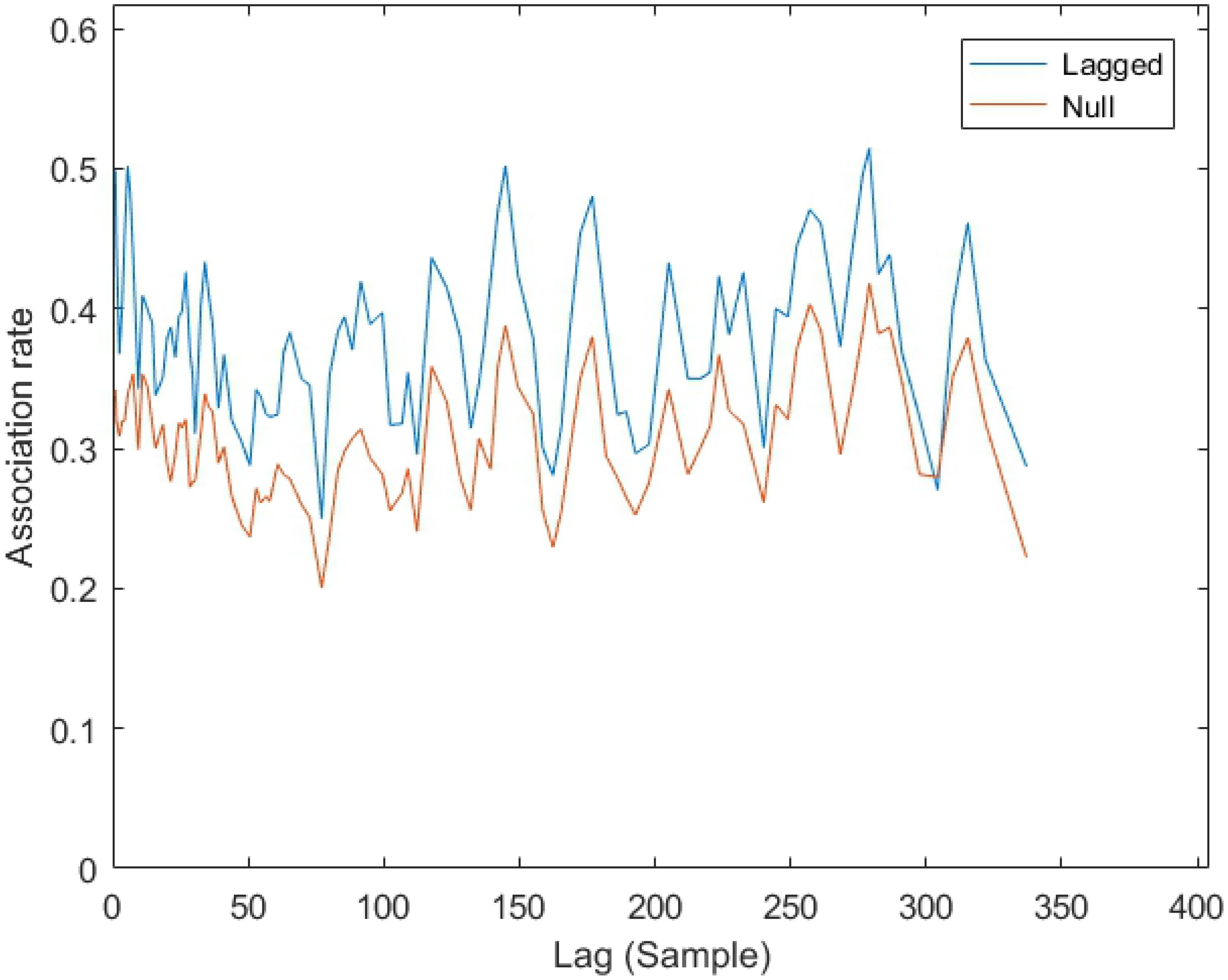
**Lagged Core Unit Association Rates** from 2-hour sample to 2-hour sample plotted relative to null association rates, demonstrating that there was a tendency for core units to stay in association and move together over long periods of time.

Hierarchical cluster analyses showed that association patterns among core units of TR band were not random but rather that some core units within the band associated preferentially, making up what we refer to as a ‘clan’. We selected the terminology from hamadryas baboon multi-level societies, where clans comprise one-male units that travel together preferentially throughout the day^24^. Association indices, where a value of 0 means that core units were never observed in association on a scan and a value of 1 means that they were always together, ranged from 0.0108 for the core unit dyad AN-BR to 0.1206 for the core unit dyad MA-PS (Fig 3). Using the dendrogram created with the average linkage method, which had the best fit with the data (CCC = 0.812), the histogram of cumulative bifurcations (Fig 4) showed one slope change that was significant at the knot AI = 0.05 (Z-score: *Z* = 1.831, *p* = 0.034). If this knot is used as a cut off to determine the number of clans present, two different clans are evident (Fig, 5 and 6), one containing three core units (FA, PS, MA) and one containing nine (PO, AN, NE, BR, FU, PH, LO, LI, AL). Permutation tests confirmed that observed association indices differed from random patterns of association^62^. Some units associated preferentially, as shown by the observed CV of association being significantly higher than in a random data set created with 10,000 permutations (CV_Obs_ = 0.484, CV_Rand_ = 0.44, *p* = 0.014). Additionally, some units showed avoidance of one another, in that the observed proportion of non-zero association indices was significantly lower than in the permuted random data set (Prop_Obs_ = 0.897, Prop_Rand_ = 0.96, *p* = 0.008).

**Fig 3.**
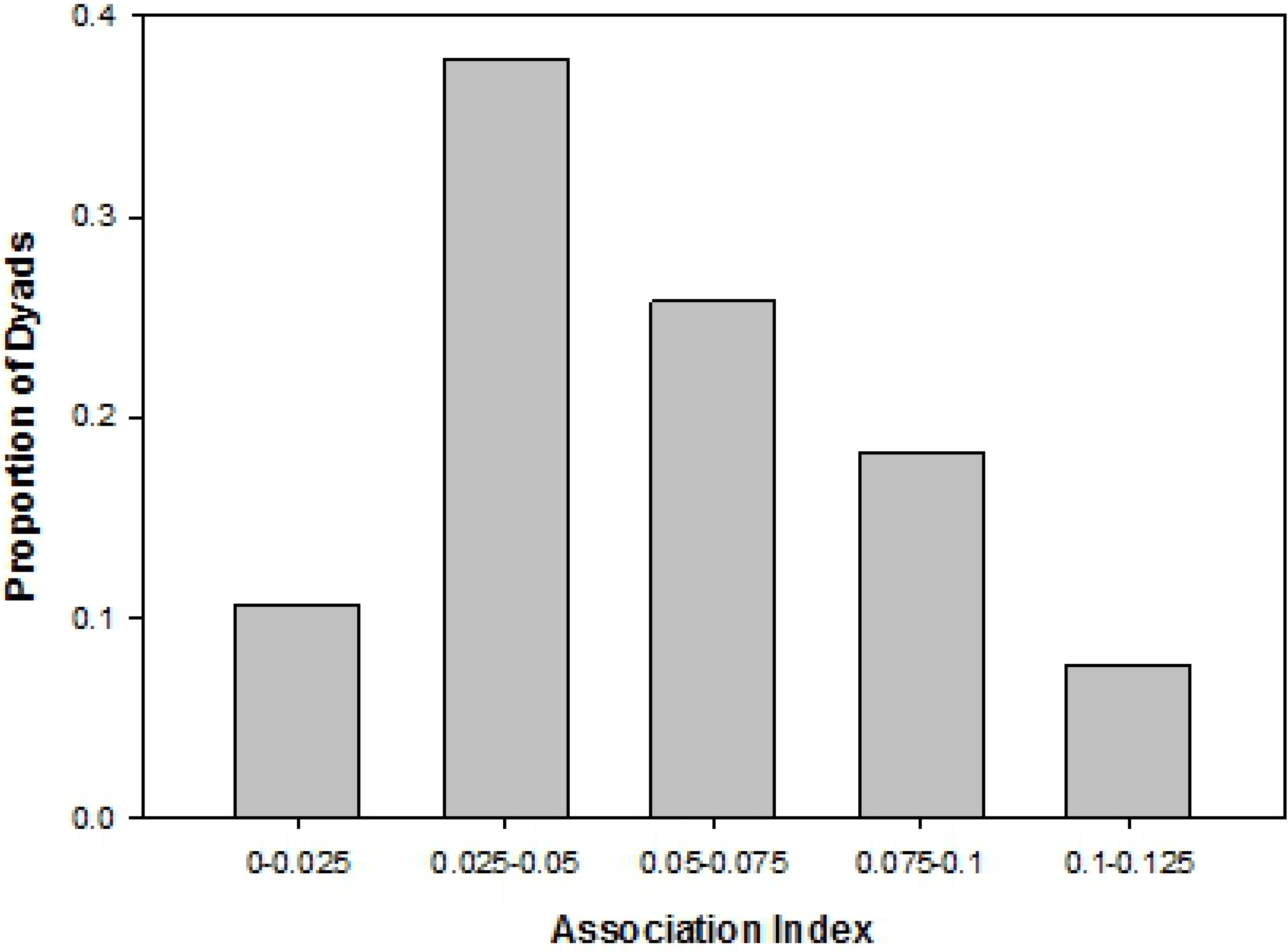
**Histogram of Association Indices** for the proportion of 66 unit dyads (12 core units) within TR band in each labeled bin. Association indices ranged from 0.0108 to 0.1206.

**Fig 4.**
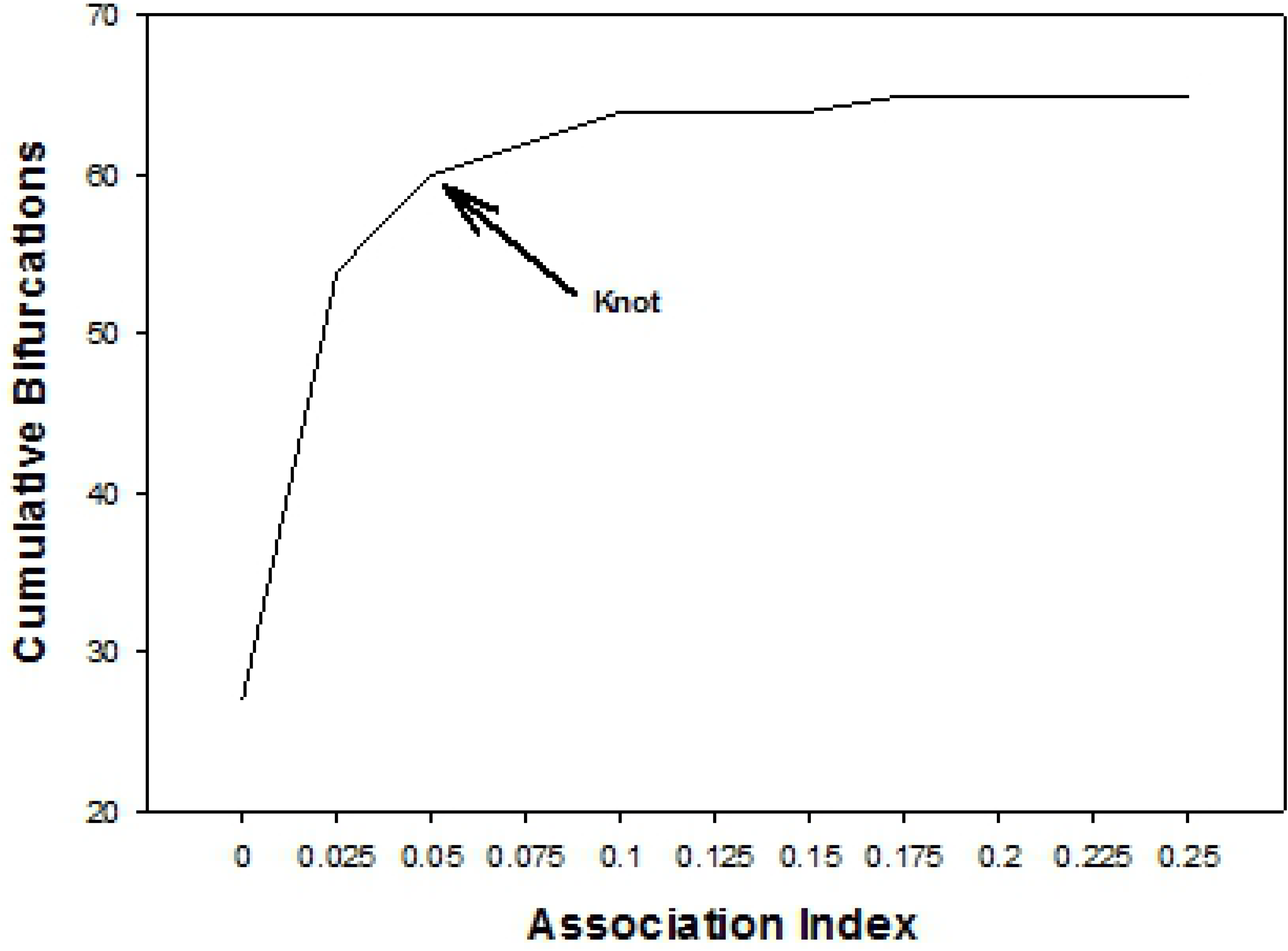
**Dendrogram of Core Unit Clusters** obtained with hierarchical cluster analyses based on average linkage. Dashed line shows that at least two clans can be differentiated at an AI of 0.05.

**Fig 5.**
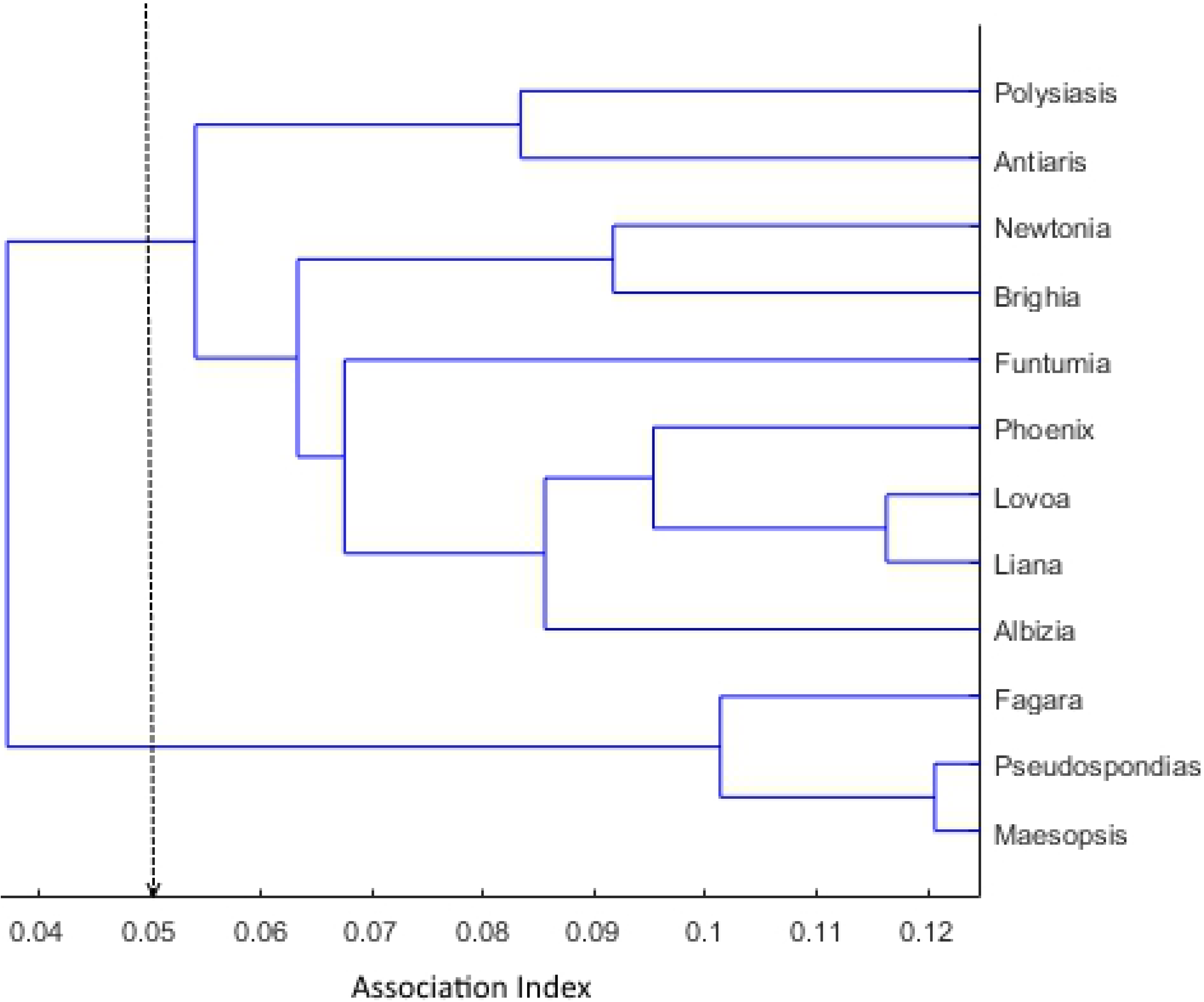
**Graph of Cumulative Bifurcations** at each increasing level of association indices for the dendrogram created using the average linkage method on core unit associations. The arrow indicates where a significant change in slope occurs (AI = 0.05), indicating a new tier of association.

**Fig 6.**
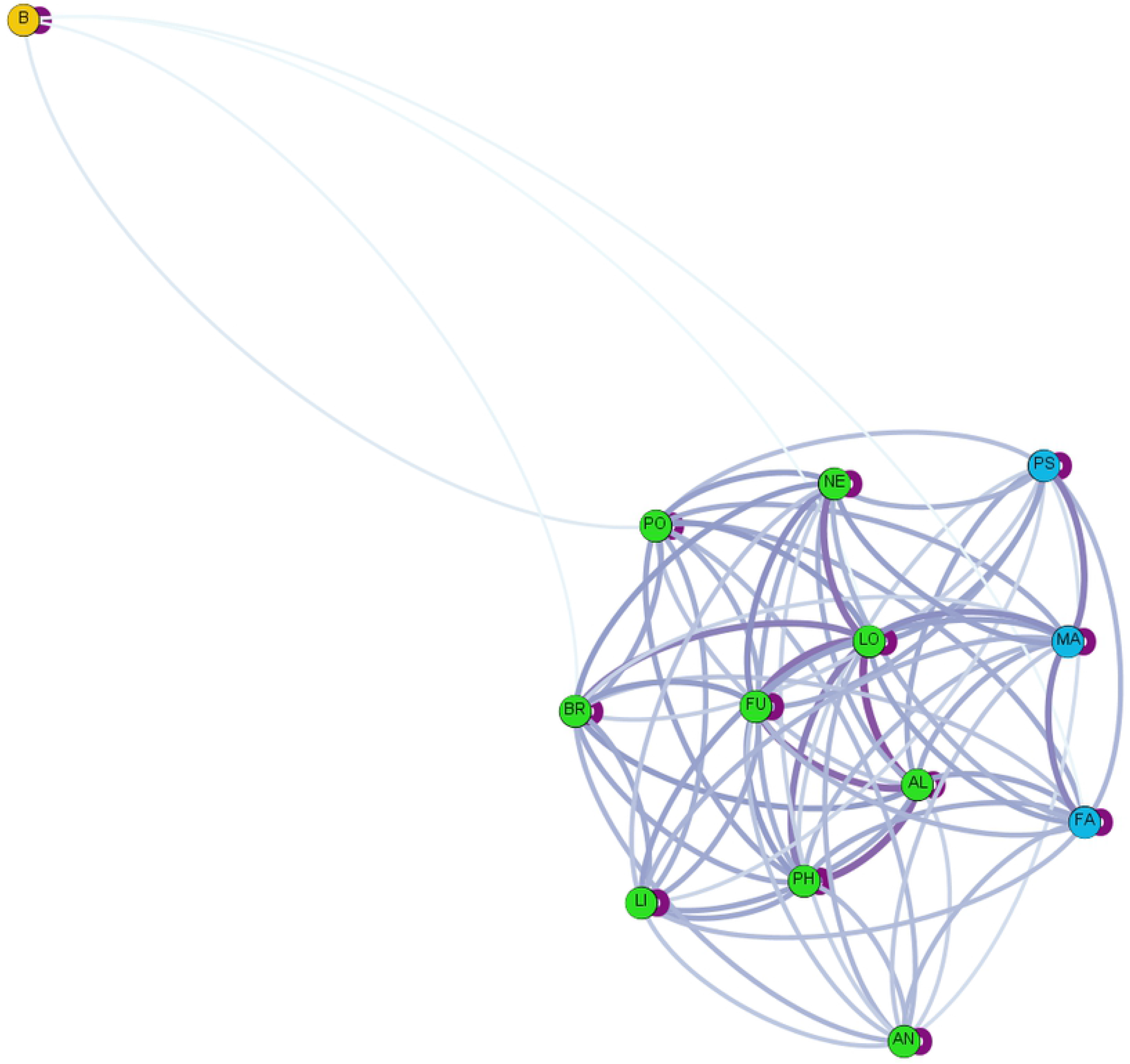
**Core Unit Social Network** based on association indices between the 12 core units in TR band and neighbouring band(s) indicated by the “B”. The darkness of the lines indicates the strength of the association. Color similarity indicates units that were determined to be in the same clan using hierarchical cluster analyses.

### Dispersal patterns

Core units had stable memberships, with transfers between units being rare. From August 2017 to August 2018, eight individuals were observed to transfer to and/or from core units within the TR band: four females and four males (Table 2). Two adult females transferred in parallel from another band and immigrated into TR band, joining core unit FU. In addition, another adult female immigrated into TR band from another band and joined core unit AN. One female transferred within TR band from core unit PS to core unit AN. Two males transferred in parallel within TR band from core unit FU to core unit PH. Finally, two additional transfers of adult males occurred within TR band, with one male transferring from core unit PH to core unit LO and one male transferring from core unit NE to core unit BR. In total, three individuals transferred to TR band from another band, all of which were female. All male transfers occurred within TR band. One additional female disappeared from her core unit and although she was seemingly healthy, we suspect that she may have died because she still had a juvenile offspring that nursed occasionally. Note that in all cases where individuals were observed to leave a core unit, transfers appeared to be voluntary.

**Table 2.** Dispersal Data for *C. a. ruwenzorii* at Nabugabo, Uganda.

| Individual | Core Unit Left | Core Unit Joined | Month |
| --- | --- | --- | --- |
| Adult Female | Another band | AN | September 2017 |
| Adult Female | PS | AN | November 2017 |
| Adult Female | Another band | FU | June 2018 |
| Adult Female | Another band | FU | June 2018 |
| Adult Male | FU | PH | July 2018 |
| Adult Male | FU | PH | July 2018 |
| Adult Male | NE | BR | June 2018 |
| Adult Male | PH | LO | May 2018 |

Table 2 shows the dispersal events that were documented from August 2017 until August 2018.

No AMUs existed during the year of data presented here, or to our knowledge, in our previous observations which were intensive by 2015. However, following our year of data collection and during the writing of this manuscript, the largest core unit (LO), which consisted of 8 adult males and 6 adult females, split permanently into two units: a OMU and a 7-member AMU. Band membership was stable throughout this study period: from August 2017 to August 2018, band TR comprised the 12 aforementioned core units. We do not currently know how clans vary in their composition over time, but this is something that we plan to investigate in the future.

## Discussion

### Delineated tiers of non-random association

This study provides the first evidence of Rwenzori Angolan colobus living in a multi-level society with three tiers of non-random association (core unit, clan, band). Our data provide evidence of 12 core units, collectively referred to as TR band, that associate non-randomly throughout their shared home range. Despite their large home range of over 1.5 km^2^, the 12 study core units were in association with at least one other core unit from TR band, on average, more than 75% of the time. There were no features within the home range where core units seemed to spend a disproportionate amount of time (i.e., rare food source, sleep sites), discounting the possibility that a spatial feature could be driving these high association rates. This suggests that they have some sort of affinity to one another. Due to the dense forest, we were rarely able to observe all the core units at once; however, on several occasions, we did see all 12 core units travelling together. Within TR band, certain core units associate more frequently than others, making up what we have referred to as clans. Two different clans were delineated, one containing three core units and one containing nine.

### Function of tiers

More data are needed to determine the function of each tier of non-random association. There is an extensive body of research on multi-level societies in arboreal Asian colobines (*Rhinopithecus* spp., *Nasalis larvatus*) that we can draw on^22–23,35,38,63–72^. It has been suggested that evenly distributed food resources allow for large aggregations in Asian colobine multi-level societies due to a decrease in feeding competition^21,43^. It is notable that the fission and fusion of core units is frequent in the lowland forest at Nabugabo but appears to be far less frequent for the large aggregations of Rwenzori Angolan colobus in the montane park at Nyungwe, Rwanda^48–50^. In Nyungwe, mature leaves and lichens are the predominant food sources^48^ but at Nabugabo, young leaves and fruit make up most of the diet, and are less widely available than mature leaves (unpubl. data). This implicates fluctuations in food availability as important in determining grouping patterns. In hamadryas baboons, it has been suggested that the clan tier evolved to allow individuals to fine-tune their group sizes to reflect local food availability^24^; a similar explanation may exist for Rwenzori Angolan colobus.

The ‘bachelor threat’ hypothesis is the most widely cited for Asian colobines and posits that ancestral, spatially-distinct OMUs aggregated so that resident males in OMUs could form coalitions in response to the threat of bachelor males^42^. Indeed, males in OMUs of Asian colobines have been observed to coordinate patrols and group defense in response to bachelor males^38,44^. Our observations thus far do not suggest that bachelor males pose a threat in Rwenzori Angolan colobus and so, we must turn to other selective forces, such as predator defense^39–40^. While native terrestrial predators appear to have been extirpated at Nabugabo, dogs kept by people pose a threat, as well as snakes and birds of prey. These hypotheses and other selective forces in the evolution of a Rwenzori Angolan colobus multi-level society require further assessment. Future research should focus on how fluctuating selective pressures (food availability, predation threat etc.) drive changes in the grouping behaviour of Rwenzori Angolan colobus at Nabugabo.

### Core unit composition

Two evolutionary pathways have been proposed for primate multi-level societies^29,34,43^. For African papionin species, phylogenetic reconstructions suggest that ancestral species had large multi-male/multi-female groups that underwent fissioning into present-day OMUs^29^. Fissioning is suggested to have occurred in response to ecological changes in the distribution of feeding sites (small and dispersed feeding patches) paired with male monopolization of females^29^. A different evolutionary route has been suggested for multi-level societies in Asian colobine species^43^. It is suggested that social pressures (i.e., bachelor males) selected for the aggregation of ancestral, spatially-distinct OMUs and that widely available food allowed for the formation of these large groups^21,43^. The core units of all primate multi-level societies studied to date contain only one reproductive male. The MMUs observed in our population of Rwenzori Angolan colobus are thus unique within the Primate Order in having several reproductive males in approximately half of the core units of the study band. Extant, closely-related species of African colobines live in either one-male/multi-female or multi-male/multi-female groups. Thus, it is possible that the evolutionary ancestors of *C. a. ruwenzorii* did as well, allowing for the aggregation of both types of core unit into a multi-level society.

### Social structure and dispersal regimes

Multi-level societies, by definition, comprise individuals and social units that associate in a non-random, hierarchical manner. This is the one unifying characteristic of all multi-level societies across the Animal Kingdom; however, in terms of the underlying social dynamics, there are several species-specific differences. This is exemplified by hamadryas baboons and geladas, two cercopithecine species that display very different dispersal regimes and social structures despite being closely related and both living in multi-level societies. In geladas, males leave their natal unit to takeover an established OMU or to join an AMU, either within the same band or another band. Female geladas remain in their natal unit, which allows for strong bonds to form amongst female kin^10–11,73^. Conversely, in hamadryas, it is the males that are philopatric while females are the dispersing sex. Females are usually coerced by males to transfer from one OMU to another^29,74^. Thus, hamadryas females do not form and maintain bonds with one another in the same way as gelada females. Instead, female hamadryas are most strongly bonded to the male in their core units (cross-bonding)^11,75–76^. While removal of a gelada adult male does not disrupt the cohesion and stability of the core unit, in hamadryas, a core unit that has lost its adult male will become unstable and the females will not remain cohesive^77–78^.

Most research on multi-level societies in Asian colobines comes from golden snub-nosed monkeys (*Rhinopithecus roxellana*, reviewed by Qi and colleagues^38^), although there is also some research on *R. bieti* ^22,35^. Published work shows that female-female bonds are critical for social cohesion while males are more socially peripheral^63–65^. This aligns with what is known about their dispersal patterns, as males tend to disperse from their natal band prior to sexual maturity, limiting the opportunities for this sex to form long-term bonds with other core unit members^66–68^ (although see ^68–69^). Previous work on our study population found that cross-sex bonding is stronger than bonding among the sexes^52^, a pattern associated with bi-sexual dispersal in other primates^80^. Bi-sexual dispersal is supported in this subspecies, as over the course of the year, we observed a total of eight dispersals, four females and four males. Interestingly, while studies on golden snub-nosed monkeys show that females transfer between OMUs of the same band, whereas males tend to disperse between bands^63,66,69^, preliminary evidence in our population of Rwenzori Angolan colobus suggests the opposite; we only observed females transferring between study bands. Of the three individuals known to have transferred to our band from another band during our study period, all were female. Each male transfer that occurred (*N* = 4) involved movement to and from core units within our study band. More data are needed to understand these patterns.

### Congeners

Indeed, it still remains a possibility that other subspecies of Angolan colobus also form multi-tiered social organizations since the six other subspecies in this species group are not well-studied. Published data on other subspecies of Angolan colobus (*Colobus angolensis* spp.) show cohesive, uni-male/multi-female and multi-male/multi-female groups of 2-20 individuals that occupy partly overlapping home ranges (reviewed by Fashing^51^). However, at Diani Beach, Kenya, *C. a. palliata* groups that range from 2-13 individuals with 1-2 males, multiple females, and offspring^81^ have been reported to form temporary associations with conspecific groups that can last from hours to days^82^. This “super-trooping” behaviour was reported to occur on 33% of observation days^82^. Larger, multi-male groups have been observed for *C. a. cottoni* in the Ituri Forest, D. R. Congo; groups of 17-20 individuals with a range of 2-5 males have been reported^83^. These groups sometimes formed “super-troops” with conspecific groups and that this occurred on 40% of observation days^82^. Though these numbers are lower than the 98% of days that we found Rwenzori Angolan colobus core units in association, further research on these, and other, subspecies of Angolan colobus could reveal that multi-level societies are in fact more widespread than currently thought in the African colobines.

## Conclusions

The discovery of another extant, nonhuman primate showing a multi-level society with female dispersal beyond the band is exciting. Like hamadryas baboons^29^, this may make *C. a. ruwenzorii* important in offering insight into human evolution. The fact that multiple reproductive males reside within a core unit in these colobus and are not overtly aggressive to one another is also of interest. Male bonding and female exogamy were key adaptations that occurred sometime during human evolution^84,85^ and thus explaining the selective pressures that led to these changes is a primary goal in anthropology.

## Acknowledgements

The authors would like to thank Colin Chapman, Lauren Chapman, Dennis Twinomugisha, Noah Snyder-Mackler, Edward Mujjuzi and Hannington Kakeeto for their help both in and out of the field.

## Supplementary Information

**S1 Dataset. Core Unit Association Dataset**

**S2 Dataset. GPS Points Dataset**

